# Combined omics unravels the molecular mechanism of golden-leaf coloration in *Koelreuteria paniculata* ‘jinye’

**DOI:** 10.1101/2022.05.19.492690

**Authors:** Ting Guo, Ruqian Wu, Xiong Yang, Sai Huang, Deyu Miao, Tingting Chen, Yinxuan Xue, Juan Li, Kai Gao, Bin Guo, Xinmin An

## Abstract

*Koelreuteria paniculata* is widely distributed in Asia and introduced to Europe and North America. *K. paniculata* ‘jinye’ is a mutant variety used in landscaping that has a golden leaf color phenotype. Although similar leaf color variants occur in plants, little is known of the underlying mechanism. We performed physiological, anatomical, microRNA sequencing, transcriptomic, and metabolomic analyses of the golden leaf variation in the mutant. Compared with the original green cultivar, the golden leaf mutant exhibited 76.05% and 44.32% decreased chlorophyll *a* (Chl *a*) and chlorophyll *b* (Chl *b*) contents, respectively, and significantly increased carotenoid content. Analysis of leaf ultrastructure revealed an abnormal chloroplast morphology and fewer lamellae in the mutant. Fifty-nine differentially expressed genes (DEGs), forty transcription factors (TFs) and forty-nine differentially expressed miRNAs (DEmiRs) involved in pigment metabolism, chloroplast development, and photosynthesis were identified. The *GLK* and *petC* genes were downregulated and are involved in chloroplast development and chlorophyll synthesis, respectively. The upregulated *PSY* and *PDS* genes, and the downregulated *NCED* gene promote carotenoid accumulation. A variety of chalcones and flavonols were upregulated in the mutant. Consequently, the carotenoid to chlorophyll ratio increased by more than 75%, and the accumulation of chalcones and flavonols was responsible for the golden leaf phenotype of the mutant *K. paniculata*.

## Introduction

Leaf color mutants are widely used for landscaping. They have bright colors, long viewing cycle, can be used to form large color blocks, and replace flowers with leaves in the light flower season. Therefore, leaf color mutant plants—such as *Ulmus pumila* ‘Jinye’, *Acer rubum, Cotinus coggygria, Populus deltoids* ‘Quanhong’, and *Photinia fraseri*—are preferred for landscaping and road greening (Zhang *et al*., 2017). Although the genetic patterns of colored-leaf plants are varied, at the metabolic level, the leaf color of higher plants depends on the content and ratio of pigments (mainly including chlorophyll, carotene, and flavonoids) in leaves. Chlorophyll is an important photosynthetic pigment, and mutations in genes linked to its biosynthesis, chloroplast protein transport, or the plant pigment regulation pathway (Wen *et al*., 2016; Zhang *et al*., 2017) can result in leaf coloration mutants. Plant carotenoids range in color from yellow to red, are involved in light harvesting, and are indispensable for photoprotection against excess light. An increase in the carotenoid-to-chlorophyll ratio may result in yellow leaf traits, such as the yellow-striped leaves of a *Ginkgo biloba* L mutant (Li *et al*., 2018). Anthocyanins are naturally occurring pigments responsible for the red, purple, and blue color of plant organs, and are essential in multiple biological processes. The formation of red leaves is the result of anthocyanin accumulation (Li *et al*., 2015). Therefore, studies of the molecular mechanism of leaf color mutants have focused on pigment metabolism.

*Koelreuteria paniculata* is an arbor species widely distributed in Asia and introduced to Europe and North America. This species is strongly adaptable to the environment and suitable for phytoremediation in heavy metal-contaminated areas (Tian *et al*., 2009). Its crude extract has medicinal and antimicrobial properties, and its leaves are used as antifungal and antibacterial agents by local people (Yang *et al*., 2018). Because of its rich flavonoids, *K. paniculata* is used in both medicine and landscaping (Kim *et al*., 2017; Lyu *et al*., 2017). *K. paniculata* ‘jinye’ is a new variety of *K. paniculata* bred by seedling mutation. Its physiological characteristics are similar to those of *K. paniculata* except that the golden leaves remain yellow during the growing period, enhancing its ornamental value. It also serves as a source of materials for study of the molecular mechanism and secondary metabolites of the *K. paniculata* leaf-color mutation.

The integration of metabolomics and transcriptomics can reveal the biosynthetic mechanisms of key differential functional pathways in plants (Li *et al*., 2020). For example, metabolomics and transcriptomics revealed the metabolic and transcriptional differences between the ‘Rougui’ protogreen leaf variety and its yellow leaf mutant, deepening our understanding of the mechanism of tea leaf coloration (Wang *et al*., 2020). Investigation of changes in the transcriptome and metabolome of jujube peel at various maturity stages and the mechanism of jujube peel coloring revealed the metabolic pathways and genes linked to jujube peel color change (Zhang *et al*., 2020). Indeed, combined -omics methods are now widely used in plant research.

To gain insight into the biological basis of leaf color and metabolite changes, we performed metabolomic, RNA-seq, microRNA sequencing and iso-seq analyses of the physiological and transcriptomic characteristics of golden leaf coloration in *K. paniculata*. Our findings provide reference information for studies of leaf coloration in other plant species.

## Results

### Cytological changes in mutant leaves

We compared chloroplast ultrastructure in normal and mutant leaves. In healthy plants, the chloroplast was oblong or fusiform, close to the cell membrane, the stromal lamella and the basal lamella were closely arranged, and the lamellae were stacked neatly. There were a few osmophilic granules and starch granules on the surface, and the chloroplast membrane was intact (Fig. 1a, b). In mutant leaves, chloroplast morphology was abnormal, the thylakoid structure was disrupted, and the grana lamella was loose and broken or missing (Fig. 1d). Correspondingly, some chloroplasts contained irregularly arranged vesicles. Furthermore, there were fewer starch granules in chloroplasts of GL, but these were filled with numerous plastoglobuli (Fig. 1c), indicating a low photosynthetic capacity and a disrupted chloroplast membrane.

**Figure 1.**
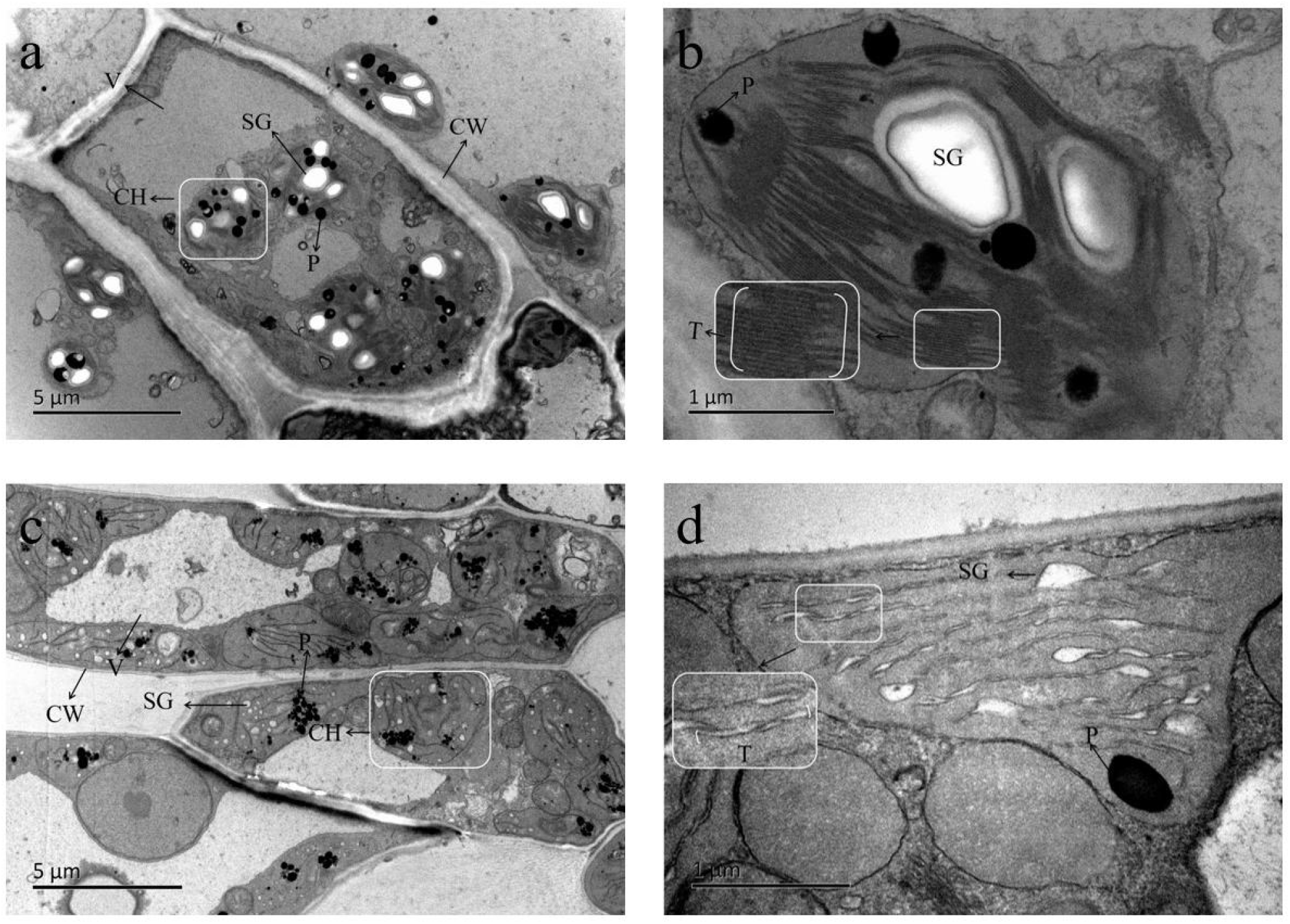
Chloroplast ultrastructure on *K. paniculata* and *K. paniculate* ‘Jinye’ in July. (a, b) The chloroplast ultrastructure of normal green leaves showed typical structure and distinct thylakoid membrane. (c, d) Abnormal chloroplast ultrastructures contained irregularly arranged vesicles in the mutant. Bars = 5µm (a, c), 1µm (b, d). CH chloroplast, CW cell wall, V vacuole, P plastoglobuli, T thylakoid grana, SG starch granule.

### Physiological changes in GL leaves

We analyzed changes in the pigment contents of WT and GL (Fig. 2). Compared with the WT, the contents of Chl *a* and Chl *b* in leaves of GL in May decreased by 87.38% and 44.19%, respectively; the total chlorophyll content decreased by 72.81%, and the anthocyanin content decreased by 10%. The content of carotenoids increased by 26.43%; the carotenoid-chlorophyll ratio in the WT was 0.35, which was significantly lower than in the mutant leaves (1.63). In July, the Chl *a* and Chl *b* contents in *K. paniculata* ‘jinye’ leaves decreased by 76.87% and 44.32%, respectively, compared with the WT whereas the contents of carotenoids increased by 30.22%, and the contents of anthocyanins remained unchanged. The carotenoid-chlorophyll ratio in GL increased by 78.49%. Compared with the WT, the Chl *a*, Chl *b*, and anthocyanin contents in GL decreased significantly in September, whereas the carotenoid content and carotenoid-chlorophyll ratio increased significantly. In May and September, there was no significant difference in the contents of chlorophyll intermediates between WT and mutant leaves (Fig. S1), but in July, the Urogen III and Pchlide contents increased significantly, whereas that of coprogen III decreased significantly.

**Figure 2.**
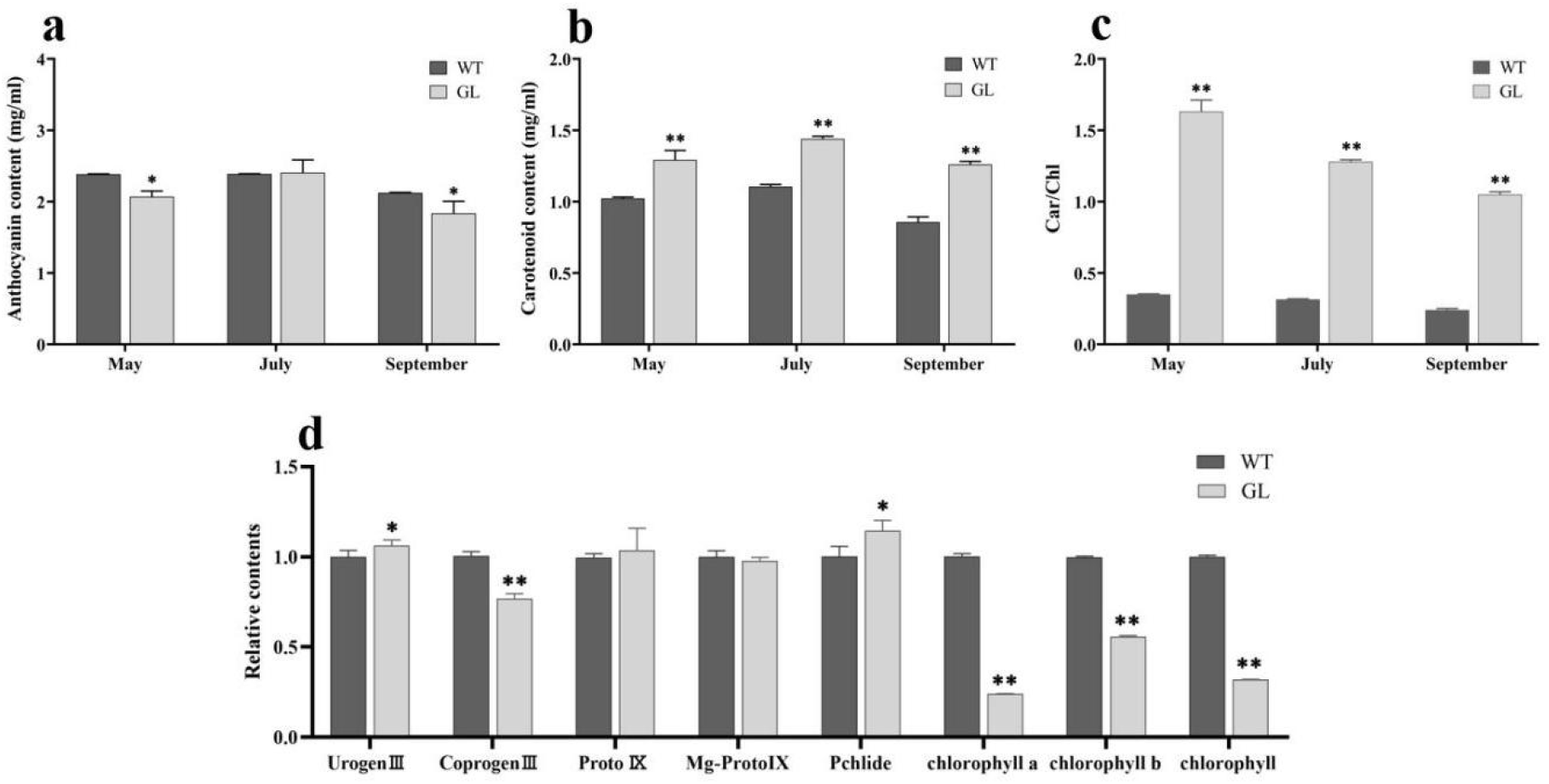
Determination of pigment contents in *K. paniculata* and *K. paniculata* ‘jinye’. (a) anthocyanin contents of normal green and mutant leaves. (b) Carotenoid contents of normal green and mutant leaves. (c) The ratio of carotenoid to chlorophyll content of normal green and mutant leaves. (d) Relative content of chlorophyll and chlorophyll intermediaries between normal green and mutant leaves in July. Asterisks indicate: (*) P⩽ 0.05, (**) P⩽ 0.01

Using LC-MS with PCA and PLS-DA analyses (Fig. S2), we detected a total of 1471 metabolites. Using a VIP ≥ 1 and fold change ≥ 2 or fold change ≤ 0.5 as criteria, 128 significantly changed metabolites (SCMs) were detected and grouped into 11 classes (Table S2). Among them, the abundances of 95 and 33 SCMs were increased and decreased, respectively. Organic acids, lipids, and oxygenous organic compounds were major contributors to the separation between the two samples. These are likely related to the basic growth metabolism of plants caused by leaf color changes. However, phenylpropane and polyketides were upregulated in GL (Fig. S3). To assess their contribution to leaf color variation, we generated a clustering heat map of 128 SCMs using the TBtools method (Fig. 3). The GL contained higher levels of flavonoids. Notably, the top enriched KEGG terms among the SCMs detected were phenylpropanoid biosynthesis, glutathione metabolism, and flavone and flavonol biosynthesis (Fig. S4). Also, chalcone, an important intermediate of flavonoid metabolites, and several flavonols, but not anthocyanins, accumulated significantly.

**Figure 3.**
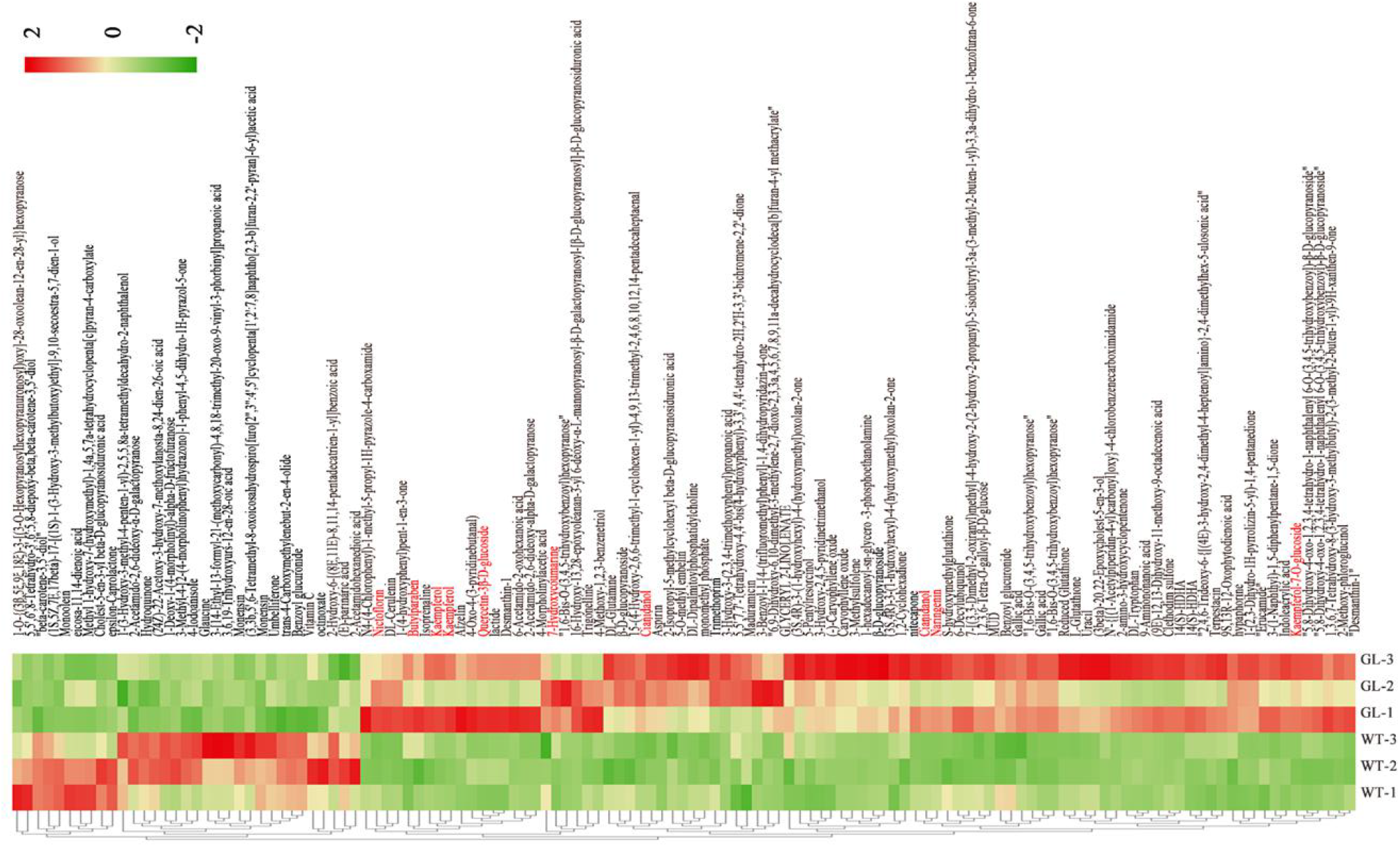
Hierarchical clustering of metabolites in WT and GL. Intensity values were adjusted by log transformation and then normalized. Flavonoid with significant differences in abundance among differently colored leaves are indicated in red text.

### DEmiRs related to pigment metabolism and predicted target genes in WT and GL

Most DEmiRs and their predicted targets were primarily identified between WT and GL in pigment metabolism pathway. Further, the correlation between DEmiRs and DEGs was analyzed. A total of 77 relevant miRNA-mRNA interaction pairs, including 45 in the chlorophyll metabolic pathway (Fig. 4a), 14 in the carotenoid metabolic pathway (Fig. 4b), and 18 in the flavonoid synthesis pathway (Fig. 4c), were predicted. Further, analyses were carried out to identify whether these interactions were either coherent (the expression level of target mRNA is more when that of miRNA is less; the “DU” and “UD” patterns) or non-coherent (miRNA and its target mRNA have similar expression profiles) (Li *et al*., 2022).

**Figure 4.**
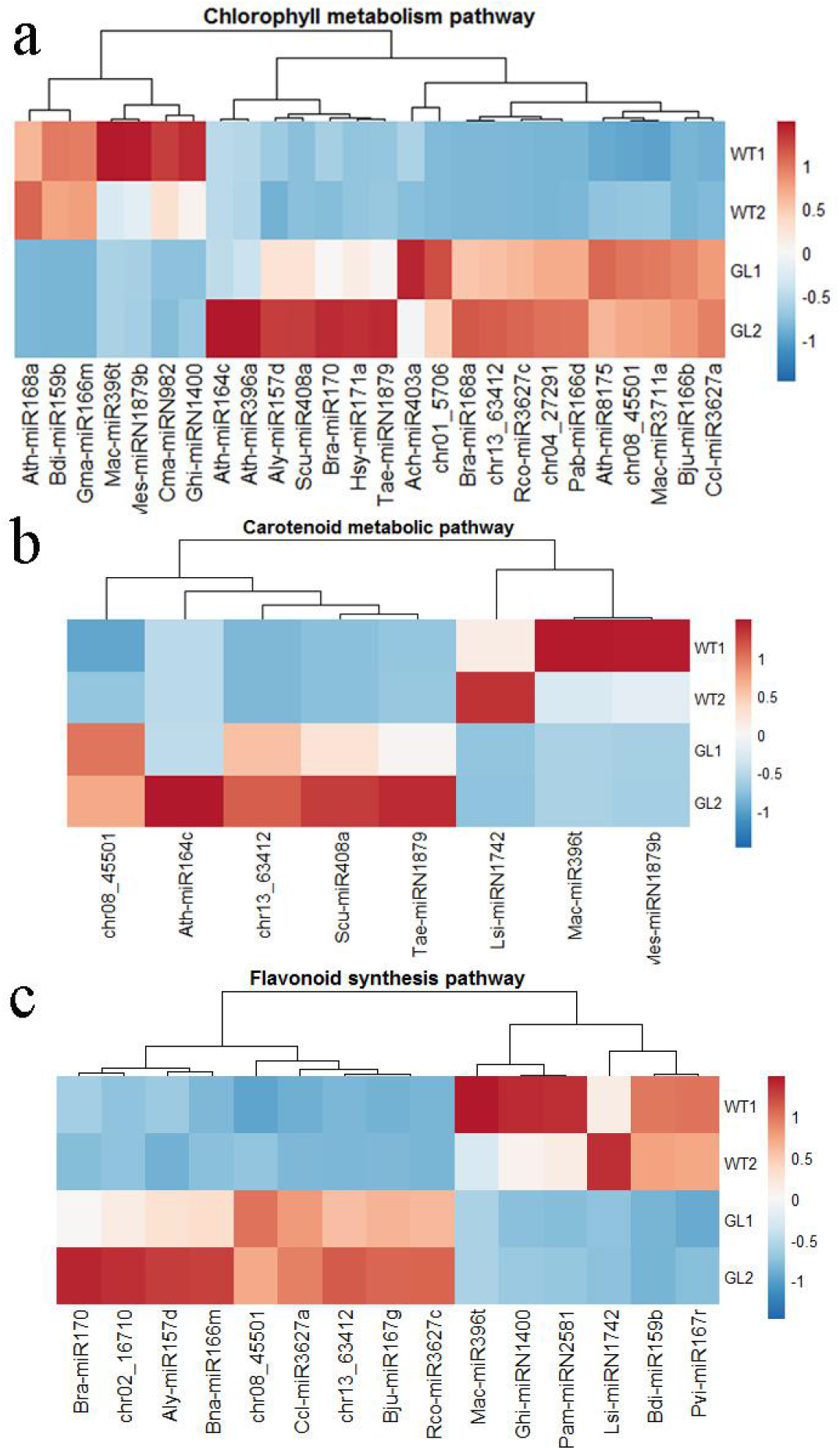
Differentially expressed miRNAs in the pigment metabolic pathway.

Analyses revealed that 77 miRNA-mRNA pairs, involving 49 miRNAs and 27 targets, were associated expressed with transcriptome data between WT and GL (Table S5, S6, S7). In most combinations, the higher the target gene expression, the lower the microRNA expression, for example miR166b-GLK, miR3711a-petC pairs etc.; a few combinations indicated similar microRNAs and target genes expression, for example miR157d-psaO etc..

### Overview of SMRT and Illumina sequencing

Using SMRT sequencing technology, 20 Gb of subreads were detected in the roots, stems, leaves, flowers, and seeds of *K. paniculata*. By filtering based on a length > 50 bp, full passes ≥ 0, and quality > 0.80, we screened 355,606 RoIs (Fig. S5), including 93.81% (333,598) FLNC reads and 3.80% (13,512) NFL reads (Table 1). The FLNC sequences were clustered to obtain nonredundant isoforms for use as a reference for sequence alignment. The clustering sequence statistics are shown in Table S3.

**Table 1.**
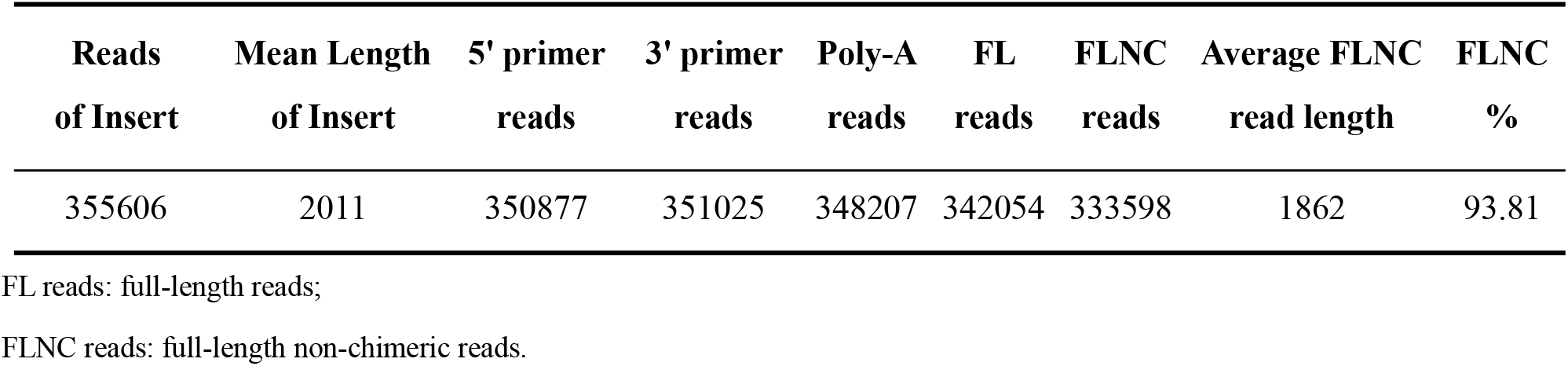
Full length transcriptome sequencing results.

From WT and GL leaves, we obtained 38.34–41.38 million raw reads, which yielded 35.57–38.32 million clean reads after quality control. The Q30 of the raw reads ranged from 89.1% to 91.69%, indicating the high quality of the transcriptome data (Table 2). Furthermore, 79,393 unigenes were annotated in six databases (Fig. 5b, c, d; Fig. S6). In the NR database, 75,344 annotations were obtained, accounting for 94.9%; the minimum number of annotations in the GO database was 26,954, accounting for 33.95%; 13.73% of the unigenes were annotated in all databases (Table S4).

**Table 2.**
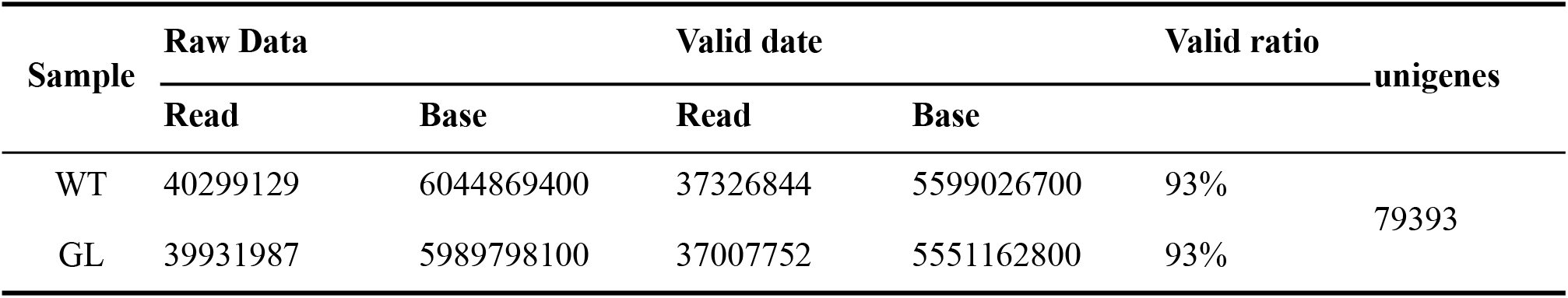
Summary of reads results before and after processing.

**Figure 5.**
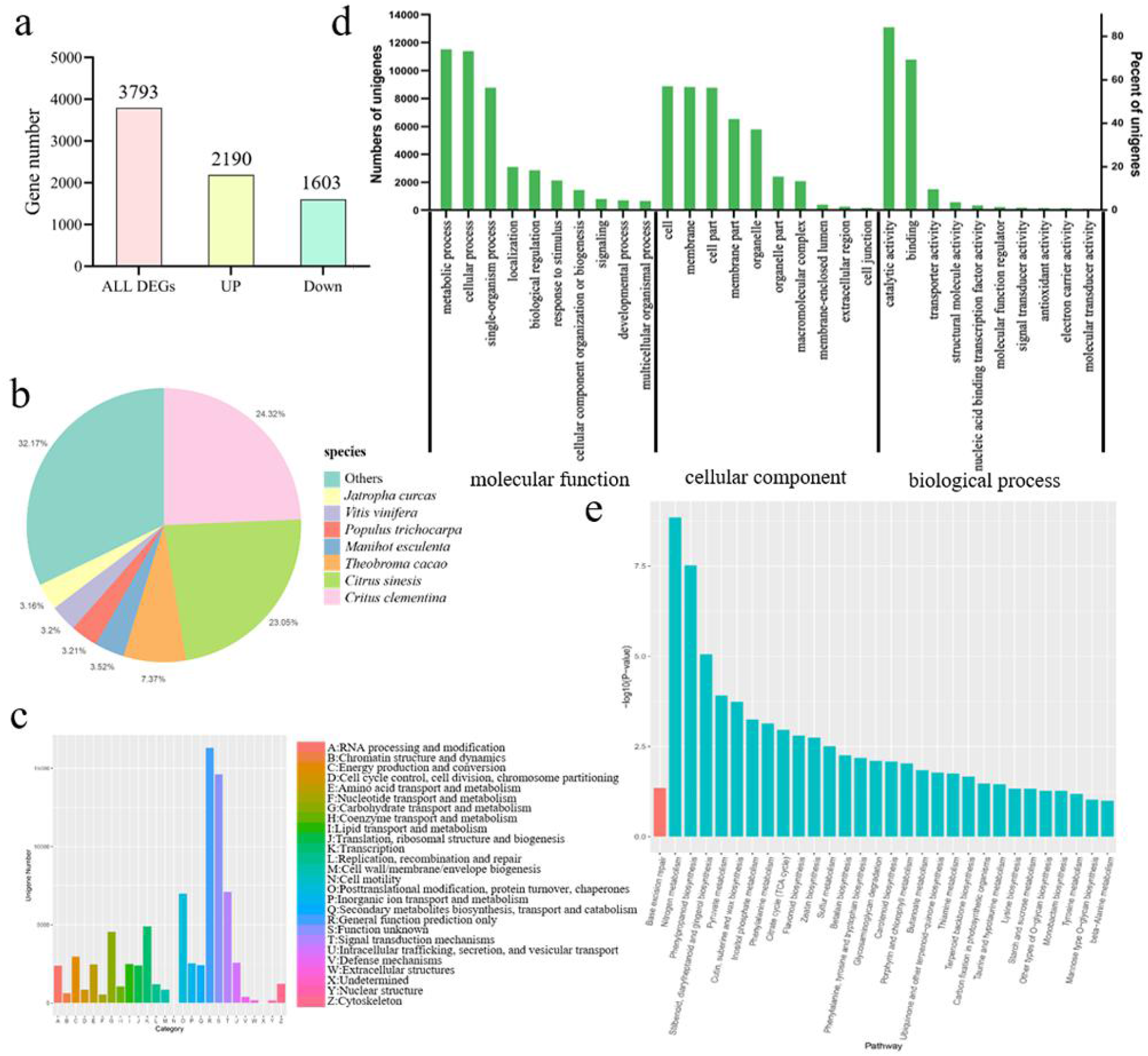
Functional annotation of unigenes in leaf transcriptomes of *K. paniculata* among different samples. (a) Summary of the transcriptome DEGs. (b) Top 7 species distribution of *K. paniculata* unigenes. (c) KOG classification of K. paniculate unigenes. (d) GO classification of K. paniculata unigenes. (e) KEGG pathway enrichment of DEGs.

### Identification and verification of DEGs

A differential expression analysis yielded 3793 DEGs in the leaves of WT compared to GL, including 2190 upregulated and 1630 downregulated genes (Fig. 5a).

GO analysis indicated that among biological processes, “carbohydrate metabolic process” and “inositol biosynthetic process” were the top enriched terms, whereas among cellular components, most DEGs were enriched in “cell wall” and “external encapsulating structure.” Among molecular functions, most DEGs were enriched in “catalytic activity” and “inositol-3-phosphate synthase activity” (Fig. S7). We speculate that a change in leaf color affects photosynthetic efficiency and alters physiological metabolism. “Organic acid metabolic process”, “L-phenylalanine metabolic process”, and “tetrapyrrole binding” were the major leaf color-related GO terms across all GO categories. Among these, tetrapyrrole binding is related to chlorophyll synthesis. “Organic acid metabolic process” and “L-phenylalanine metabolic process” are upstream pathways of flavonoid synthesis. In short, in response to leaf color mutation, there may be a variety of changes in biological response in *K. paniculata*.

The KEGG pathway analysis showed a significant separation between the WT and GL, indicating that changes in metabolite accumulation during development are tightly governed by differential gene expression. The WT *vs*. GL DEGs mapped to 121 KEGG pathways. Among them, primary metabolic processes were significantly enriched, demonstrating that leaf-color changes affect plant growth and development. Six pathways related to leaf color were identified: “phenylpropane biosynthesis”, “flavonoid biosynthesis”, “porphyrin and chlorophyll metabolism”, “carotenoid biosynthesis”, “photosynthesis”, and “photosynthetic antenna protein”. Of the top KEGG enrichment pathways, two were related to flavonoid metabolism (“phenylpropanoid biosynthesis” and “flavonoid biosynthesis”) (Fig. 5e). In mutant leaves, many downregulated DEGs were enriched in “energy metabolism” and “carbohydrate metabolism” whereas several upregulated DEGs were enriched in “porphyrin and chlorophyll metabolism” and “carotenoid biosynthesis.” A few DEGs were enriched in the photosynthesis pathway.

### DEGs and SCMs related to pigment metabolism in WT and GL

Normal green leaves depend on the balance of chlorophyll, carotenoids, and flavonoids. Although the content of Coprogen III in yellow leaves was about 0.77-fold that of green leaves, the contents of other intermediates such as UrogenIII, Proto IX, Mg-ProtoIX, and Pchlide were not decreased in yellow leaves. This is consistent with the finding that 17 DEGs annotated in the porphyrin and chlorophyll metabolism pathway were upregulated in mutant leaves (Fig. 6a). However, the Chl *a* and Chl *b* contents were lower in yellow leaves compared to green leaves. RNA-seq showed that the expression of *CHLG*, which catalyzes chlorophyllide production, was not significantly affected in yellow leaves. *GLK* (golden2-like) genes are key regulators of chloroplast development. In this study, the expression of two DEGs annotated as *GLKs* was lower in yellow leaves. According to the KEGG pathway results, 15 unigenes in the photosynthesis pathway were annotated as 10 genes. Compared with normal green leaves, the expression levels of *psaA, psaF*, and *psaO* (related to the reaction center subunit of photosystem I [PSI]), as well as *ATPB* (related to F-ATPase) were upregulated. However, the mRNA levels of *ATPD, ATPG*, and *petC* (associated with the cytochrome b6-f complex) were significantly downregulated, and *petC* was a highly significantly downregulated.

**Figure 6.**
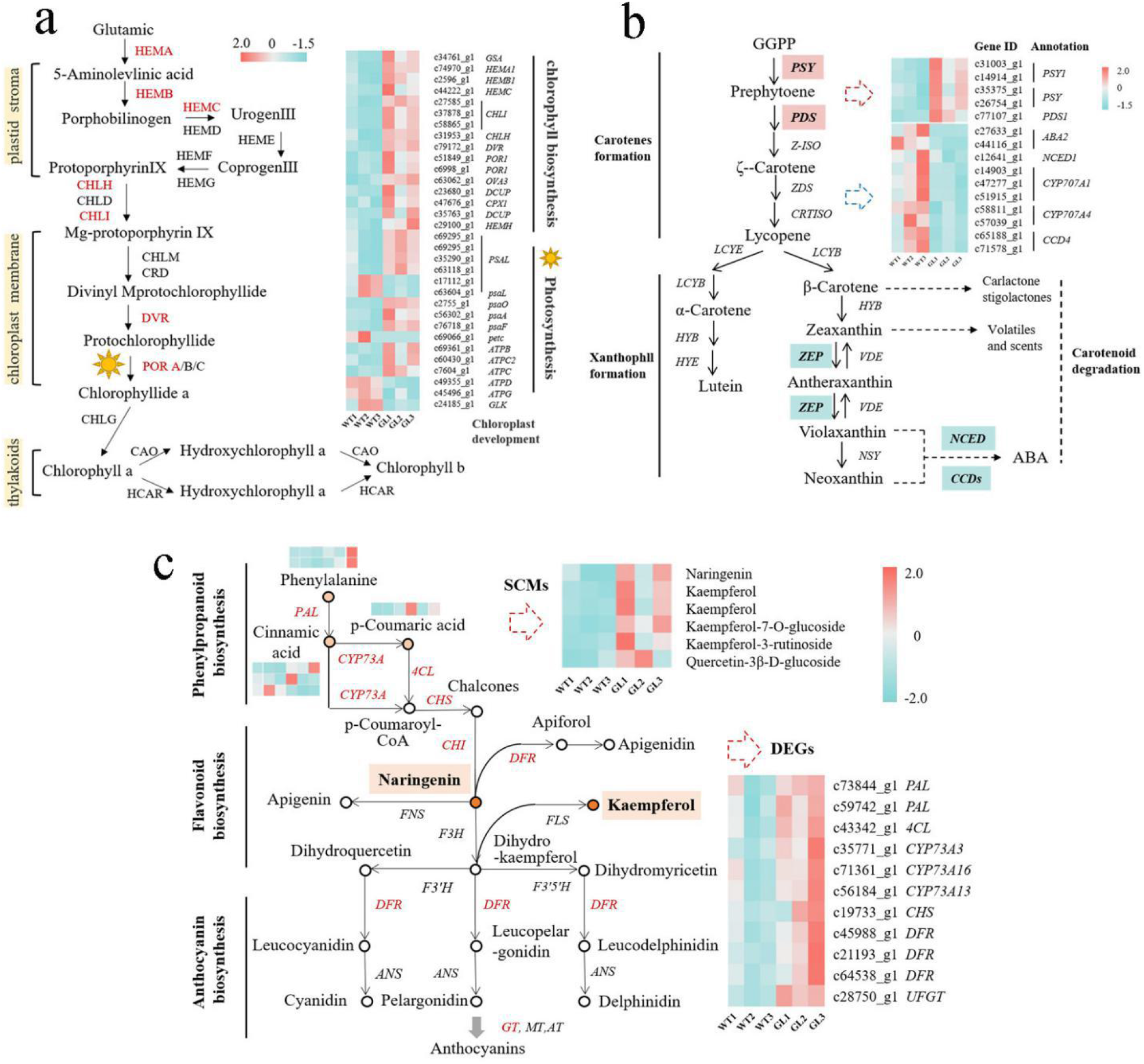
Differentially expressed genes (a) Chlorophyll metabolic pathway (b) Carotenoid metabolic pathway (c) Flavonoid synthesis pathway

The total carotenoid content differed significantly between WT and GL, and carotenoid composition may influence leaf coloration. We identified 15 DEGs regulating carotenoid biosynthesis and degradation based on KEGG pathway annotations (Fig. 6b). Two genes involved in carotenoid metabolism were upregulated in mutant leaves; one was annotated as PHYTOENE SYNTHASE (*PSY*) and the other as 15-CIS-PHYTOENE DESATURASE (*PDS*). Nine carotenoid cleavage dioxygenase (*CCD*) genes encode synthetic proteases, which mediate oxidative degradation of carotenoids. Three *CCD1/CCD4/NCED* genes, which are closely related to carotenoid degradation, were significantly downregulated in the mutant. *ABA2, CYP707A1*, and *CYP707A4* were related to the biosynthesis of abscisic acid (ABA) downstream of the carotenoid synthesis pathway, and all were downregulated. The upregulation of carotenoid biosynthesis genes and downregulation of carotenoid degradation genes likely lead to carotenoid accumulation in mutant leaves.

Based on the KEGG enrichment results, 2 of the top 10 metabolic pathways were related to flavonoid synthesis. We constructed a network to assess the relationships between gene expression and metabolite accumulation (Fig. 6c). Compared with the WT, the abundance of cinnamic acid and *p*-coumaric acid in GL was increased. The intermediates naringenin and kaempferol, as well as other flavonoids and flavonols, accumulated significantly in mutant leaves. We identified 17 DEGs associated with flavonoid biosynthesis. The expression of nine DEGs was upregulated in GL leaves, including flavonoid synthesis precursor synthase genes (*e*.*g*., *PAL, 4CL*, and *CYP73A*), early biosynthesis genes (*e*.*g*., CHS, CHI), and a late biosynthesis gene anthocyanin biosynthesis gene (*DFR*). This is consistent with the higher flavonoid and flavonol contents in the leaves of GL.

### Transcription factors involved in leaf coloration

In this study, 381 DEGs were identified as transcription factors (TFs) belonging to 39 TF families, some of which were significantly upregulated and downregulated. Therefore, leaf-color variation affects the regulation of gene expression via TFs. The most abundant TF family was the MYB ultrasound family (53, 13.91%), followed by the C3H (36, 9.45%), NAC (30, 7.87%), ERF (27, 7.09%), bHLH (19, 4.99%), and WRKY (17, 4.46%) families (Fig. S9). MYB and bHLH transcription factors regulate flavonoid biosynthesis. In this study, the MYB family was represented by 43 DEGs upregulated in GL. Also 19 DEGs related to bHLH transcription factors were identified, most of which were upregulated in GL. In addition, the expression of DEGs encoding WRKY, TCP, and C2H2 family TFs was significantly higher in GL leaves than in WT leaves.

### Verification of DEGs by RT-qPCR

To verify the transcriptome data, we performed RT-qPCR analysis of selected DEGs at three developmental stages (May, July, and September). In total, 15 DEGs were detected, comprising 5 genes involved in chlorophyll biosynthesis, 2 in chlorophyll development, 2 in photosynthesis, 3 in carotenoid metabolism, and 3 genes involved in flavonoid biosynthesis. The expression levels of 12 genes detected by RT-qPCR showed patterns similar to the transcriptome data (Fig. S10). In May, July, and September, the expression levels of several genes in the mutant were significantly decreased, such as *GLK* and *FtsH* (related to chloroplast development) and *NCED* (related to carotenoid degradation) (Fig. S10h, i, l). *PDS* and *CYP73A* were upregulated in the mutant (Fig. S10k, n). Compared with the WT, *HEMA* (involved in chlorophyll biosynthesis) was significantly upregulated in GL in May and September but downregulated in July (Fig. S10a). *HEMC* was significantly upregulated in May and downregulated in July and September (Fig. S10b). In May and July, photosynthesis-related genes were significantly downregulated, but significantly upregulated in September (Fig. S10f). By contrast, *psaL* was significantly upregulated in May and July and downregulated in September (Fig. S10g). *PSY* and *DFR* were consistent with the trend of *psaL* (Fig. S10j, o). In general, the RT-qPCR results were consistent with RNA-seq data, indicating the transcriptome data to be valid and reliable.

## Discussion

We investigated a yellow leaf mutant of *K. paniculata*. To explore mechanisms of the leaf color mutation in *K. paniculata* ‘jinye’, numerous color-related genes were identified and their expression patterns were investigated. We focused on the genes related to pigment synthesis, chloroplast development, TFs, and photosynthesis based on enrichment results of transcriptome and metabolome analyses.

### Chlorophyll metabolism and chloroplast development are responsible for leaf color change

Chlorophyll is an important pigment in the thylakoid that captures and transfers light energy in photosynthesis (Nakajima *et al*., 2012). Leaf color formation is closely related to chlorophyll metabolism and chloroplast development. Similar to other yellow-leaf mutants, *K. paniculata* ‘jinye’ is a chlorophyll-deficient chlorina mutant. Compared with the WT, the Chl *a* and Chl *b* contents of GL were significantly lower by 76.05% and 44.32%, respectively. Therefore, the decrease in total chlorophyll content was responsible for the yellow leaf phenotype.

In higher plants, the synthesis of chlorophyll starts from glutamyl tRNA, and can be divided into three parts: the synthesis of 5-aminolevulinic acid (ALA) to protoporphyrin IX is completed in the chloroplast matrix, and the synthesis of magnesium protoporphyrin to chlorophyllide is completed in the chloroplast membrane. Finally, Chl *a* and Chl *b* are synthesized on the thylakoid membrane (Matringe *et al*., 1992). Most leaf color mutants have an altered thylakoid membrane structure. For example, in yellow leaf mutant *G. biloba*, chloroplast ultrastructure was markedly altered, chloroplast-like membranes were broken, vesicles were dense, and there were no inner membrane structures (Li *et al*., 2018). Similarly, an abnormal internal capsule membrane was found in *P. deltoides*, bamboo, and *Anthurium andraeanum* leaf mutants (Yang *et al*., 2015). In this study, the chloroplast structure of GL differed significantly from that of the WT. The chloroplasts of mutant leaves were irregularly shapes, in the basic structure, the thylakoid membranes were ruptured and chloroplast grana were loosely arranged and showed fractures and deletions. Also, there were more plastoglobuli and fewer starch particles; the latter indicates decreased photosynthetic capacity, and osmium-containing granules indicate disruption of the chloroplast membrane. These structural abnormalities in chloroplasts affect photosynthesis and chlorophyll accumulation.

In this study, precursors of chlorophyll synthesis did not decrease in the mutant, and no gene in the chlorophyll biosynthesis pathway showed decreased expression. As an important TF for chlorophyll accumulation and chloroplast formation, *GLK* was significantly decreased in GL, similar to reports of other species (Gang *et al*., 2019). For example, low expression of *dmGLK2* and no expression of *dmGLK1* inhibited chloroplast development and decreased the chlorophyll content, resulting in the yellow-green leaf phenotype of chrysanthemum (Chang, 2013). A 40-bp deletion mutation in *BpGLK1* of birch decreased the chlorophyll content, hampered chloroplast development, and induced a yellow-green leaf mutant (Gang *et al*., 2019). Therefore, a change in chlorophyll content may be reflected by abnormalities of plastid development and function.

### Photosynthesis is responsible for the leaf-color change

Light is the most important environmental factor affecting plant leaf color (Zhang *et al*., 2019). The leaf color of *K. paniculata* ‘jinye’ was yellow-green in the shade and after shading treatment, the leaf color changed from golden yellow to yellow-green. This suggests *K. paniculata* ‘jinye’ to be a photosensitive mutant. The leaf color of the GL mutant is affected by light intensity (Zhang *et al*., 2019). However, the mechanism of the light response of the GL mutant was unclear. Light is necessary for chloroplast morphogenesis. Light controls gene transcription, chlorophyll synthesis, and protein degradation, thus regulating chloroplast development. The phenotype of chlorophyll-deficient mutants was affected significantly by environmental factors such as light (Song *et al*., 2017). During photosynthesis, light energy is captured by the pigment in the LHC protein and transferred to the reaction center complex of photosystem I (PSI) and photosystem II (PSII). The cytochrome b6-f (Cyt b6-f) complex responds to the electron flow balance from PSII to PSI via the plastid quinone pool and regulates the activity of PSII kinase light-harvesting complex. In a chlorophyll-deficient mutant, LHC protein was significantly reduced or lost, hampering granule accumulation in chloroplasts (Kim *et al*., 2009). In this study, the expression of three DEGs in the LHC gene family was decreased in GL, indicating that the light-harvesting chlorophyll protein was reduced compared with the reaction center complex. PSI catalyzes light-driven electron transfer from luminosome cyanin to matrix ferredoxin, which consists of more than 10 subunits including PsaC and PsaL. The PsaL subunit is responsible for most of the interactions, and its mutants show a slightly smaller functional size of the photosynthetic antenna and a lower excitation level (Klodawska *et al*., 2013). Activity of the Cyt b6-f complex, which is encoded by chloroplast and nuclear genes, affects the electron transfer rate. The Cyt b6-f complex Rieske Fe/S precursor protein of all photosynthetic eukaryotes is encoded by the nuclear *petC* gene (Veit *et al*., 2012). Site-directed knockout of the *petC* gene in *Arabidopsis thaliana* blocks the electron transport chain and produces a yellow leaf trait (Maiwald *et al*., 2003). In a *Forsythia* GL mutant, *petC* expression was downregulated and not affected by light intensity (Shen, 2019). In this study, *psaL* and *petC* expression in the Cyt b6-f complex decreased highly significantly (log2foldchange –9.1). Therefore, downregulation of gene expression in photosynthesis may explain the decreased chlorophyll content. *petC* (c69066_g1) is key for the leaf color change in *K. paniculata* ‘jinye’.

### Carotenoid biosynthesis is responsible for the leaf color change

Carotenoids have important effects on photomorphogenesis and photosynthesis (Cazzonelli and Pogson, 2010; Li *et al*., 2009). Carotenoids have multiple conjugated double bonds and can absorb light in the range of 400–500 nm. Therefore, the accumulation of carotenoids turns plants yellow, orange, and red (Zhou *et al*., 2020). Compared with the WT, the carotenoid content in GL leaves increased significantly. Yellow leaf mutants typically have a significantly increased carotenoid content. Carotenoid accumulation and pigmentation are dependent on the expression of carotenoid biosynthesis genes (Hao *et al*., 2020). As the first committed enzyme in carotenoid synthesis, phytoene synthase (PSY) converts two molecules of geranylgeranyl pyrophosphate (GGPP) to phytoene (Kang *et al*., 2014). Overexpression of *PSY1* increased the carotenoid content of tomato fruit (Fraser *et al*., 2007). PDS catalyzes the biosynthesis of lycopene from phytoene (Fu *et al*., 2016). In this study, five DEGs (*PSY, PSY1*, and *PDS*) related to carotenoid biosynthesis were upregulated in GL and likely contribute to carotenoid accumulation.

Carotenoid accumulation is also related to carotenoid degradation. Plants have two major carotenoid degradation pathways mediated by carotenoid dioxygenase (CCD) and 9-*cis*-epoxidized carotenoid dioxygenase (NCED). The degradation of carotenoid is suppressed in CCD1 mutant Arabidopsis, the seed of which showed an increased carotenoid content (Schwartz, 2001). Moreover, the β-carotene content of the flesh of a *CCD4*-deficient mutant changed at the late-ripening stage, accompanied by yellowing (Ma *et al*., 2014). *NCED* is the rate limiting step from 9’-*cis* neoxanthin to ABA. The transcriptional regulation of *NCED* is a focus of research on ABA metabolism. RNAi-mediated inhibition of *NCED* expression in tomato increased the lycopene and β-carotene contents of mature fruit and reduced the ABA content (Ji *et al*., 2014). The mRNA levels of other ABA synthesis-related genes, such as *ABA2* and *CYP707A*, were significantly decreased in GL. As a downstream product of the carotenoid pathway, ABA may be a feedback regulator of carotenoid metabolism (Mohd *et al*., 2013). Exogenous ABA induces important genes involved in carotenoid metabolism, such as *PSY3* in maize and *DSM2*, a *β-Carotene Hydroxylase* gene in rice (Du *et al*., 2010; Li *et al*., 2008). Moreover, some TFs in citrus, such as CrMYB68, indirectly affect carotenoid metabolism by directly inhibiting ABA biosynthesis. The effect of ABA on carotenoid metabolism needs further study.

### Flavonoid biosynthesis is responsible for leaf color changes

Flavonoids are an important part of leaf color formation and protect leaves from damage caused by sunlight-derived UV irradiation (Agati *et al*., 2012). In this study, GO and KEGG analyses identified 17 DEGs in flavonoid metabolic pathways and 10 flavonoid-related DEGs in phenylpropanoid biosynthesis. Most showed significant changes in expression level. The transcript abundances of flavonoid synthesis precursor synthases (*e*.*g*., PAL, 4CL, and CYP73A), EBGs (*e*.*g*., CHS and FLS), and LBGs (*e*.*g*., DFR) were higher in GL compared to WT. Phenylalanine ammonia lyase (PAL) catalyzes the deamination of L-phenylalanine (L-Phe) to yield cinnamic acid as the first committed step in phenylpropanoid synthesis. *4CL* encodes 4-coumaric acid coenzyme A ligase, which catalyzes the conversion of phenylalanine to 4-coumaric acid coenzyme A as a precursor for flavonoid synthesis. As key enzymes in flavonoid biosynthesis, PAL and 4CL affect the total flavonoid pathway flux through a variety of phenylacetone pathways (Winkel-Shirley, 2001). The upregulation of upstream gene expression may lead to accumulation of reaction substrates (Liu *et al*., 2019). High expression of EBGs leads to the accumulation of chalcones and flavonols. Although DFR was upregulated, no anthocyanins were found among SCMs. This may be a result of redirection of carbon flux to flavonoid branches, which is consistent with the significant increase in kaempferol content in the GL.

As secondary metabolites of phenylpropanes, flavonoids have a basic structure of C6-C3-C6 and are classified into several groups (*e*.*g*., chalcones, flavonols, flavones, and anthocyanins). Flavonoids have the widest color range, from light yellow to blue. Anthocyanins are responsible for the plant coloration range from orange to blue and are used as natural food pigments. Chalcones and aurones are the main sources of yellow color in plants. For example, naringenin is widely found in yellow plants, whereas aurones are endemic to *Caryophylla* (Tanaka *et al*., 2008). Some flavonols and flavones are light yellow, affecting anthocyanin color development as co-pigments. In this study, most differential metabolites were chalcones, flavonols, and flavones, but not anthocyanins, which explains the yellow color of the leaves. Our results shed light on the regulatory network of flavonoid biosynthesis. The combination of metabolomics and transcriptomics enables investigation of the relationships between key genes and metabolites in biosynthetic pathways. We identified candidate genes and metabolites involved in the flavonoid biosynthesis pathway, providing insight into the leaf color variation of *K. paniculata*.

## Conclusions

We investigated differences in coloration between normal green leaves of *K. paniculata* and gold-color leaves of *K. paniculata* ‘jinye’ at the physiological and molecular levels. The mutant leaves showed a decreased chlorophyll-carotenoid ratio and abnormal chloroplast ultrastructure. We identified 3793 DEGs and 128 SCMs in GL compared to WT. And 49 DEmiRs were identified in chloroplast development, photosynthesis, and pigment metabolism pathways. Downregulation of genes related to chloroplast development and photosynthesis, such as *GLK* and *petC*, resulted in decreased chlorophyll accumulation and enhanced carotenoid biosynthesis, increasing the carotenoid content in mutant leaves. In addition, the high expression of phenylpropanoid and flavonoid pathway genes and the accumulation of flavonoids and flavonols in the mutant may also be related to leaf color formation. We conclude that the formation of leaf color of *K. paniculata* ‘jinye’ is related to chloroplast development, photosynthesis, and pigment metabolism pathway genes. This leads to accumulation of carotenoids, flavonoids, and flavonols, and a decrease in chlorophyll content, resulting in a change in leaf color from green to yellow (Fig. 7). Our findings provide a reference for studying the mechanism of plant leaf color variation.

**Figure 7.**
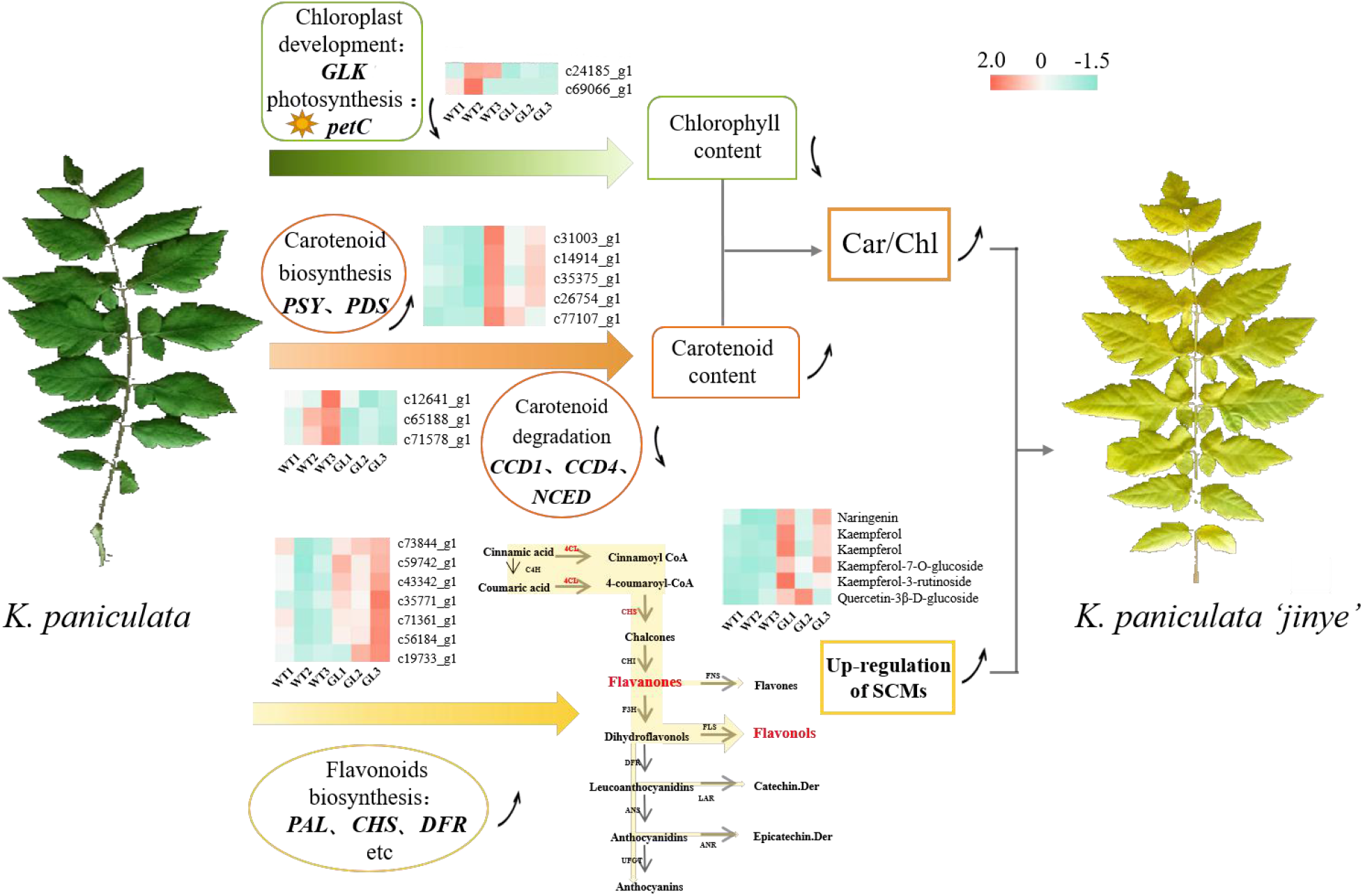
The proposed pathway of mutant leaf coloration in *K. paniculata*

## Materials and Methods

### Plant materials

The green-leaf *K. paniculata* cultivar (wild type, WT) and the yellow-leaf *K. paniculata* ‘jinye’ cultivar (golden leaf mutant, GL) were used with three independent biological replicates. The plants were grown under natural conditions in the *Koelreuteria* nursery in Xiangyuan County, Shanxi Province, China. Leaf tissues were collected from May to September in 2018. For cytological analysis, RNA-seq, and metabolomics experiments, green leaves and mutant leaves were sampled separately on July (Fig. 8a-f). For physiological and RT-qPCR experiments, mutant leaves and green leaves collected at three stages (May to September) were used. At least 10 tender leaves at the third position from the top of a branch were sampled from three plants of each type, immediately frozen in liquid nitrogen, and stored at −80°C for further analysis.

**Figure 8.**
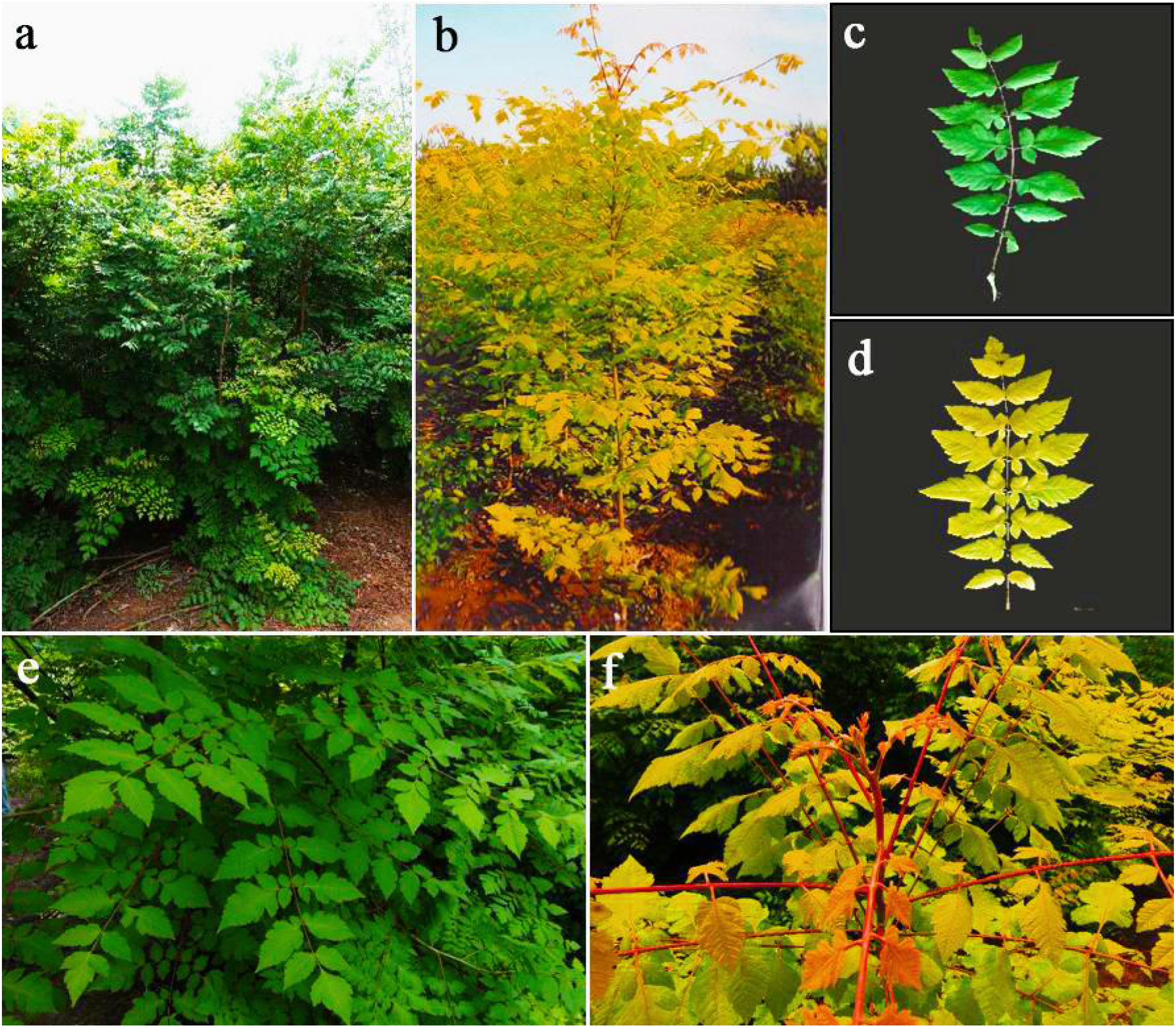
Phenotypes of leaves of *K. paniculata* and *K. paniculata* ‘Jinye’ in July. (a, c, e) Phenotype of the normal green leaves. (b, d, f) Phenotype of the mutant leaves.

### Measurement of chlorophyll and carotenoid contents

Approximately 0.1 g of wild-type and mutant leaves were cut into pieces and extracted with 15 mL of 80% acetone at 4°C for 24 h in the dark. The extract was measured spectrophotometrically at 470, 646, and 663 nm. The chlorophyll *a* (Chl *a*), chlorophyll *b* (Chl *b*), carotenoid, and cyanidin contents were determined as described elsewhere (Gang *et al*., 2019). To measure the contents of chlorophyll intermediates, leaves were dissolved in nine volumes of 0.01 M phosphate-buffered saline in an ice bath and centrifuged (30 min at 2500 × *g*). The supernatant was assayed separately using an ELISA Kit (HengYuan Biological Technology Co., Ltd, Shanghai, China). Three biological replicates were analyzed per sample. Data were analyzed by *t*-test using SPSS software (ver. 17.0; SPSS Inc., Chicago, IL), and means were compared at significance levels of 0.01 and 0.05.

### Transmission electron microscopy (TEM)

Mature leaves (third pair of leaves at the top of the plant) of the WT and GL were collected for TEM analysis. Avoiding the main vein, we cut the fresh tissue into 1–2-mm^3^ pieces and transferred them to 2.5% (v/v) glutaraldehyde for vacuum infiltration. Next, the pieces were pre-fixed in 2.5% glutaraldehyde for 24 h at 4°C, followed by 1% OsO4 for 2 h. After dehydration, infiltration, and embedding, the pieces were sectioned using an EM UC6 ultramicrotome (Leica Microsystems GmbH, Wetzlar, Germany), and observed using a JEOL 1200 transmission electron microscope (JEOL Ltd., Tokyo, Japan).

### Metabolite profiling

Metabolites were extracted from WT, mutant leaves and the equivalent mixture of two materials taken as samples. Samples (0.1 g) were analyzed by liquid chromatography electrospray ionization tandem mass spectroscopy (LC-ESI-MS/MS) with SIMCA-P data-analysis software. Next, a principal component analysis (PCA), partial least-squares discrimination analysis (PLS-DA), and an orthogonal partial least-squares discrimination analysis (OPLS-DA) were performed to verify the reliability of the data. Fold-change (FC) (FC > 2 or FC < 0.5 and P < 0.05) and variable importance for projection (VIP) (VIP > 1 and P < 0.1) criteria were applied to identify differential metabolites.

### Small RNA sequencing

#### Library preparation and sequencing

Two clones of the same cultivars in both the WT and the GL libraries were used for the small RNA sequencing. Small RNA libraries were constructed using the KAITAI-BIO Small RNA Library Prep Kit (KAITAI-BIO, China) according to the manufacturer’s instructions. The library preparations were sequenced on an Illumina HiSeq platform.

#### Data analysis

The raw reads obtained were processed. Fastp software is used to filter Raw Data, mainly including removing primer and connector sequences; low quality sequences at the end were removed (>20% bases <Q30, and N base >10%); and those shorter than 18 nt and longer than 30 nt were also discarded.

### Identification and annotation of the miRNAs

The derived reads were further screened against rRNA, tRNA, snRNA, snoRNA and other ncRNA as well as repeat sequences by mapped to RNA database, which is Rfam (http://rfam.xfam.org/) and the remaining unannotated reads were then mapped onto the *Sapindus mukorossi* reference genome.

Importantly and notably, there was not miRNA annotation in the *K. paniculata* full-length transcriptome. Therefore, to identify known miRNAs, the mapped reads with the *K. paniculata* reference genome were mapped onto the miRNAs of *Sapindus mukorossi*, which is the most evolutionarily close to *K. paniculata*.

The unaligned unique reads were further used for novel miRNA prediction using miRDeep-P2 (Kuang et al., 2018) and newly updated criteria for plant miRNA criteria (Axtell and Meyers, 2018), referring to the method described by Guo et al. (2020). Furthermore, the identified miRNAs were clustered into families based on sequence similarity.

### Differential expression analysis of miRNAs

To identify differentially expressed miRNAs (DEmiRs), edgeR was used. To filter out miRNAs with low expressions, those with |log2(FC)| ≥ 1.00 and a Benjamini-Hochberg FDR corrected P - value < 0.05 were assigned as differentially expressed. MiRNAs related to chloroplast development, photosynthesis and pigment metabolism pathway were screened from differentially expressed miRNAs, and analyzed in heat maps.

### Target genes prediction

Potential miRNA target genes were predicted using the psRNA Target, RNAhybrid Target and miranda Target. Target genes of miRNAs related to chloroplast development, photosynthesis and pigment metabolism pathway were predicted and counted.

### PacBio library preparation and sequencing

To obtain integrated full-length transcriptome sequences of *Koelreuteria*, we pooled total RNA of the root, stem, leaf, seed, and flower in equal quantities to construct sequencing libraries. Next, we used the SMARTer PCR cDNA Synthesis Kit to synthesize full-length cDNA. After end repair, adaptor ligation, and index code addition, PCR amplification was conducted. The polymerase-bound template was bound to MagBeads and sequencing was performed on a PacBio Sequel platform to obtain polymerase reads.

By filtering out the adapter sequences and subreads < 1000 bp and the raw polymerase read fragment sequences < 50 bp or sequence accuracy of < 0.80, we extracted the suitable subreads from the polymerase reads. The reads of insert (RoIs) obtained by screening were classified into full-length non-chimeric (FLNC), non-full-length (NFL), full-length-chimeric, and short (< 300 bp) reads depending on whether a 5ʹ-primer, 3ʹ-primer, or poly-A tail was detected. Finally, similar sequences among the FLNC sequences were clustered to nonredundant isoforms after removing redundant high-quality consensus FLNC reads using CD-HIT program with a threshold of 0.99 identity.

### RNA-Seq library preparation, sequencing, and expression estimation

Total RNA was isolated from the wild type and mutant with a Quick RNA Isolation Kit (Omega) and used for RNA-seq. We performed 1% agarose gel electrophoresis and used the Agilent 2100 Bioanalyzer to assess RNA integrity. RNA purity and concentration were measured using a NanoDrop 2000 spectrophotometer (Thermo Fisher). Library construction and RNA-seq were performed by HTHealth Biotechnology Co., Ltd. (Beijing, China). The libraries were subjected to next-generation sequencing on the Illumina HiSeq platform. Finally, we used in-house Perl scripts to process the low-quality data to obtain clean data.

We combined the unigenes with the PacBio Iso-Seq data resulting in a final reference transcriptome. Isoform and unigene expression levels were quantified using RSEM software (https://github.com/deweylab/rsem), and transcript lengths were normalized to FPKM values. The unigenes were mapped to five public databases: NCBI nonredundant protein sequences (NR), Swiss-Prot, Gene Ontology (GO), evolutionary genealogy of genes: Non-supervised Orthologous Groups (eggNOG), and Kyoto Encyclopedia of Genes and Genomes (KEGG), using BLAST software to obtain annotation information. We used DESeq software (Anders *et al*., 2010) to analyze gene expression with |log2foldchange| > 1 and p < 0.05 as the screening criteria. Finally, the DEGs were subjected to GO and KEGG analyses.

### Quantitative RT-qPCR

RNA-seq samples were subjected to RT-qPCR to assess the reliability of the transcriptome data. RNA extraction and cDNA synthesis were conducted as above. The 2× SYBR Green qPCR Mix Kit (TaKaRa, Japan) and the cDNA concentration and primers described above were used; the reaction volume was 25 μL and PCR was conducted using the 7500 Real-Time PCR System (Applied Biosystems). Three independent biological replicates per sample and three technical replicates per biological replicate were analyzed. The primer sequences are listed in Supplementary Table S1. DEG expression was normalized by the 2^−ΔΔCt^ method (Livak *et al*., 2001).

## Supplemental data

**Supplemental Figure S1** Relative content of chlorophyll and chlorophyll intermediaries between normal green and mutant leaves. (a) Plant leaves in May. (b) Plant leaves in September.

**Supplemental Figure S2** Score plot of metabolite profiles of J1-J3 (GL) and B1-B3 (WT).

**Supplemental Figure S3** Classification of differential metabolites

**Supplemental Figure S4** KEGG enrichment of different metabolites

**Supplemental Figure S5** Full length transcriptome sequencing data. (a) ROI sequence length distribution. (b) Pie chart of ROI classification. (c) ROI sequence quality distribution map. (d) Sequence length distribution after clustering

**Supplemental Figure S6** KEGG classification of *K. paniculata* unigenes.

**Supplemental Figure S7** GO enrichment of *K. paniculata* unigenes.

**Supplemental Figure S8** The difference expression results of metabolites and related transcripts

**Supplemental Figure S9** Distribution of transcription factors

**Supplemental Figure S10** RT-qPCR analysis of the expression of fifteen DEGs at different developmental stages between K. paniculata and K. paniculata ‘Jinye’. Asterisks indicate: (*) P⩽ 0.05, (**) P⩽ 0.01

**Supplemental Table S1** Primer sequence of RT-qPCR **Supplemental Table S2** Classification of differential metabolites **Supplemental Table S3** Statistical table of clustering sequence **Supplemental Table S4** Summary of annotation results

**Supplemental Table S5** List of miRNAs related to chlorophyll metabolic pathway and their target genes in two samples

**Supplemental Table S6** List of miRNAs related to carotenoid metabolic pathway and their target genes in two samples

**Supplemental Table S7** List of miRNAs related to flavonoid synthesis pathway and their target genes in two samples

## Acknowledgements

This work was supported by the National Natural Science Foundation of China (31870652). We thank Mr. Wang Guodong for providing experimental materials. The English in this document has been checked by at least two professional editors, both native speakers of English. For a certificate, please see: http://www.textcheck.com/certificate/te8dE6

